# A whole blood approach improves speed and accuracy when measuring mitochondrial respiration in intact avian hematocytes

**DOI:** 10.1101/2022.05.25.493398

**Authors:** Andreas Nord, Imen Chamkha, Eskil Elmér

## Abstract

Understanding mitochondrial biology and pathology is key to understanding the evolution of animal form and function. However, mitochondrial measurement often involves invasive, or even terminal, sampling, which can be difficult to reconcile in wild models or in longitudinal studies. Non-mammal vertebrates contain mitochondria in their red blood cells, which can be exploited for minimally invasive mitochondrial measurement. Several recent bird studies have measured mitochondrial function using isolated blood cells. Isolation adds time in the laboratory and might be associated with physiological complications. Inference may also be constrained on biological grounds by lack of tissue context. We developed and validated a protocol to measure mitochondrial respiration in bird whole blood. Endogenous respiration was comparable between isolated blood cells and whole blood. However, oxidative respiration was higher in whole blood, and whole blood mitochondria were better coupled and had higher maximum working capacity. Whole blood measurement was also more reproducible than measurement on isolated cells for all traits considered. Measurements were feasible over a 10-fold range of sample volumes, though both small and large volumes were associated with changes to respiratory traits. The protocol was compatible with long-term storage: after 24 h at 5 °C without agitation all respiration traits but maximum working capacity remained unchanged, the latter decreasing by 14%. Our study suggests that whole blood measurement provides faster, more reproducible, and more biologically (tissue context) and physiologically (mitochondrial integrity) relevant assessment of mitochondrial respiration. We recommend future studies to take a whole blood approach unless specific circumstances require the use of isolated blood cells.

## INTRODUCTION

The ability to convert energy contained in food into a currency that can be used to sustain bodily housekeeping functions and key fitness-related traits such as growth, reproduction, movement, and survival, is a fundamental requirement for life. This occurs by digestion of food items into macromolecules that are subsequently used to create energy for working cells in the form of adenosine triphosphate (ATP). More than 90% of ATP is produced during cellular respiration in the mitochondria (Nicholls & Ferguson 2013). It is therefore not surprising that mitochondrial respiration and/or volume increase when organismal demand for energy increase (e.g. Liknes & Swanson 2011; Zheng *et al*. 2014; Dawson *et al*. 2016). Mitochondrial ATP is created by transport of electrons through a series of protein complexes (the electron transport system, ETS) whilst protons are being actively pumped across the inner mitochondrial membrane to establish electrochemical potential. The energy required for oxidative phosphorylation (OXPHOS) is harnessed when protons backflow into the mitochondrial matrix at complex V, the ATP-synthase. However, some protons forego OXPHOS by leaking back into the mitochondrial matrix spontaneously or under the influence of uncoupling proteins (Nicholls & Ferguson 2013). Proton leak thus uncouples mitochondrial respiration from ATP production (LEAK) but generates heat and reduces the rate at which potentially biodegrading reactive oxygen species (ROS) are formed (Brand *et al*. 2004; Speakman *et al*. 2004). It follows that there is intricate regulation of the balance between LEAK and OXPHOS, such that mitochondria can be made more effective for a given substrate oxidation rate when energy is in short supply (Trzcionka *et al*. 2008; Bourguignon *et al*. 2017; Roussel *et al*. 2020), or more thermogenic to aid endothermic heat production (Walter & Seebacher 2009).

Studies of mitochondrial function can provide proximate and ultimate depth in many fundamental areas of research within ecology and evolution (see Koch *et al*. 2021 for a recent review). However, such studies are rare in comparison to the rich body of literature on organismal-level adaptation, function, and performance, particularly in the wild. This probably reflects logistic and ethical considerations, because mitochondrial measurement often requires terminal sampling for organ harvest or anaesthesia to collect biopsies, together with speedy handling of samples to ensure organelle integrity; all of which are easier to reconcile with laboratory models. However, birds and some other non-mammal vertebrates retain the nucleus and functional organelles throughout erythroblast maturation and so contain mitochondria in their erythrocytes (red blood cells) (Moritz, Lim & Wintour 1997; Moras, Lefevre & Ostuni 2017). Studies on birds have found that this makes it possible to assess mitochondrial function even from small (25-50 μl) blood samples (Stier *et al*. 2013; Stier *et al*. 2017; Stier *et al*. 2019) that can be safely collected in organisms < 5 g (Kramer & Harris 2010). This is not possible when measurements are performed using other mitochondria-rich hematocytes, as is necessary in mammals (e.g. Sjövall *et al*. 2013a; 2013b). While much is still unknown about the function and regulation of blood cell mitochondria relative those in other tissues, several recent bird studies indicate that blood cell respiration varies predictably in line with organismal-level energy demand over long (Stier *et al*. 2019; Casagrande *et al*. 2020) and short (Garcia Diaz et al. 2022) time frames, and in the contexts of seasonal adaptation (Nord *et al*. 2021) and metabolic senescence (Dawson & Salmón 2020). Moreover, blood cell mitochondrial function seems amenable to epigenetic programming during embryonic development (Udino *et al*. 2021; Stier, Monaghan & Metcalfe 2022), much like what is a well-known fact within human medicine (Gyllenhammer *et al*. 2020). Studies have even found that blood cell respiration can potentially be used to predict variation in whole-organism metabolic rate (Malkoc, Casagrande & Hau 2021). Thus, there is ample evidence that mitochondrial measurement in red blood cells can provide minimally invasive insights into several key areas within the broader fields of ecology, ecological physiology, and evolutionary biology.

To the best of our knowledge, all studies on avian blood cell mitochondria have been performed using isolated blood cells, prepared using standard laboratory techniques including pipetting, vortexing and centrifugation. These procedures could impact the status of the cells going into the experiment (e.g. Wiegmann *et al*. 2017). If standard handling impacts cell health, mitochondrial function in isolated hematocytes may not be as representative of the *in vivo* state, or could introduce experimental variation, but this notion remains untested. Furthermore, while erythrocytes will be responsible for most of the respiration in the avian circulation (c.f. Scanes 2022), it can be difficult to reliably separate cell types by centrifugation in the small samples that are typically collected from wild models. Thus, the exact composition of a cell isolate may, or may not, correspond to that in whole blood. Moreover, on conceptual grounds it can be argued that isolate-based measurements lack some biological value since the removal of plasma before measurement effectively also removes the tissue-context of the blood. On this background, we asked whether mitochondrial measurement on whole blood samples were comparable to measurements on isolated hematocytes from the same blood volume. If cell isolation impacts mitochondrial integrity or function of the electron transport chain, we predicted that isolate measurements would be associated with leakier mitochondria and/or lower overall respiration compared to whole blood samples. To explore the utility of whole blood measurement, we proceeded by testing the resilience of the assay to variation in sample volume and long-term storage. Finally, we undertook initial validation of a protocol for permeabilising whole blood to permit studies of mitochondrial function with unlimited substrate supply.

## MATERIALS AND METHODS

### Animals and housing

The study was performed using captive zebra finches (*Taeniopygia guttata* Vieillot) (*n* = 23; 12 females and 11 males). All birds were in their second calendar year, or older, and body masses ranged 12.1 to 19.4 g. The birds originated from a captive population kept in outdoor flight aviaries at the Lund University field station Stensoffa (WGS84 DD: N55.69534, E13.44739) and were brought to light- and temperature controlled indoor facilities at Lund University 5-7 d before the experiment started. Birds were kept in cages measuring 120 × 80 × 100 cm (length × depth × height) at a density of ≤ 6 birds to a cage. Room temperature was maintained at 19-20°C and photoperiod was 12:12 LD. Food (mixed millet seeds), cuttlebone, and water was provided *ad libitum*.

### Anaesthesia and blood sampling

We sterilised the ventral skin area covering the distal part of the sternum using 70% ethanol and immediately anaesthetised the birds by intraperitoneal injection of pentobarbital sodium (30-40 μg per g body mass). When the bird was reactionless to an external stimulus (toe pinching), which occurred within 6-10 min after the injection, we collected a maximal blood sample (250-650 μl) from the jugular vein and then euthanised the bird by cervical dislocation. Samples were stored at 10-15°C in 2 mL K2-EDTA (ethylenediaminetetraacetic acid) tubes (BD Vacutainer^®^, Franklin Lakes, NJ, USA) without agitation until analysed 15 to 45 min later.

### Mitochondrial respiration measurements and experiments

Mitochondrial respiration was measured at bird body temperature (41°C) (Prinzinger, Pressmar & Schleucher 1991) using Oxygraph O2k high-resolution respirometers (Oroboros Instruments, Innsbruck, Austria). Respiration was inferred from the decline in O_2_ concentration in a 2 ml suspension of sample and respirometry medium (MiR05: 0.5 mM of EGTA, 3 mM of MgCl_2_, 60 mM of K-lactobionate, 20 mM of taurine, 10 mM of KH_2_PO_4_, 20 mM of HEPES, 110 mM of sucrose, and 1 g l^−1^ free fatty acid bovine serum albumin, pH 7.1).

All measurements were performed on intact cells. We allowed 10-15 min for O_2_ consumption to stabilise after adding sample to the chambers, and then recorded baseline phosphorylating respiration on endogenous substrates for 2-4 min (“ROUTINE”). We then added 1 μg ml^−1^ oligomycin to inhibit ATP synthase, thus preventing oxidative phosphorylation. The remaining respiration in this stage is used to offset the leak of protons that occur across the inner mitochondrial membrane (“LEAK”). Thus, the part of respiration devoted to oxidative phosphorylation (“OXPHOS”) can be derived as the difference between the ROUTINE and LEAK states. This was followed by titrating the mitochondrial uncoupler carbonyl cyanide-p-trifluoro-methoxyphenyl-hydrazone (FCCP) in 0.5 μl, 1 mM, aliquots until maximum respiration was reached (final concentration: 1-2 μM). At maximal non-inhibiting concentration, FCCP abolishes the proton gradient across the inner mitochondrial membrane, forcing the electron transport system to work at its maximum capacity (“ETS”) to restore it. After FCCP titration, we inhibited mitochondrial complex I, using 2 μM rotenone, and then added 1 μg ml^−1^ of the complex III inhibitor antimycin A to stop electron transport. Any respiration remaining after addition of antimycin A is considered of non-mitochondrial origin. There was never a meaningful further reduction in respiration when antimycin A was added onto rotenone (data not shown). Hence, we adjusted all respiratory states for non-mitochondrial respiration by subtracting whichever was the lowest of respiration on rotenone and rotenone+antimycin A, under the assumption that any absolute differences between the two were due to random noise.

In the first experiment, we compared mitochondrial respiration traits measured in whole blood and isolated blood cells. Data were collected from 7 birds (3 males, 4 females) and all samples were run in simultaneous duplicates within 45 min of collection. We manually mixed the blood by gently tilting the sample tubes for 4-5 min. Then, we collected two 100 μl whole blood samples to use either for immediate respiration measurement, or for isolating the hematocytes. We isolated the blood cells following Stier et al. (2017) with slight modifications according to Dawson & Salmón (2020) and Nord et al. (2021). Thus, the sample was first centrifuged at 3000 g for 10 min at room temperature to separate plasma from the blood cells. The pellet was then re-suspended in 500 μl cold MiR05, centrifuged at 1000 g for 5 min and the supernatant was discarded. The remaining pellet, containing all cells from the 100 μl blood sample, was immediately re-suspended in 750 μl MiR05 pre-equilibrated at 41 °C and added to 1.35 ml MiR05 contained in the respiration chamber (final volume 2 ml) for use in the assay. The sample volume used here is the same as in the original protocol for mitochondrial measurement in bird blood cells (Stier *et al*. 2017) and is within range of that used in other studies (e.g. Dawson & Salmón 2020; Nord *et al*. 2021).

In the second experiment, we asked how blood sample volume affected respiration traits, attempting to refine the protocol by defining the lowest possible volume yielding representative data. Blood samples were drawn and managed as above (*n* = 7; 3 females, 4 males). Mitochondrial respiration traits were then assayed simultaneously in 10, 25, 50 and 100 μl whole blood samples. The volume of MiR05 in the chambers was adjusted accordingly to maintain a total volume of 2 mL

In the third experiment, we investigated the effect of storage time on whole blood respiration, using data from 6 birds (3 females, 3 males). Within 30 min of blood sampling (henceforth, time = ‘0 h’), we followed the protocol above to measure mitochondrial respiration in 50 μl blood samples in 1.95 mL MiR05. The tube with remaining blood was then stored without agitation at 5.2 ± 0.7 °C (mean ± s.d.; range 3.9 to 6.5 °C) and another 50 μl aliquot from the same sample was measured again (after 5 min of mixing) 24 h later (henceforth, time = ’24 h’). Samples from each of two birds were also opportunistically measured at 72 h and 96 h after collection.

### Data analyses

We calculated three flux control ratios (FCR) to address integrity of mitochondrial respiration following the method and terminology proposed by Gnaiger (2020): 1) E-R control efficiency (1 - ROUTINE / ETS), which is a measure of the proportion of maximum working capacity remaining during endogenous respiration; 2) R-L control efficiency ((ROUTINE - LEAK) / ROUTINE), which is the proportion of endogenous respiration channelled towards ATP production via oxidative phosphorylation; 3) E-L coupling efficiency (1 - LEAK / ETS), which is indicative of how ‘tightly’ electron transport is coupled to ATP production under a stimulated cellular state.

Statistical analyses were performed using R 4.1.2 for Windows (R Core Team 2021). To compare the utility of blood cells and whole blood (i.e., Experiment 1), we fitted trait- and FCR-specific linear mixed effect models (lmer in the lme4 package) (Bates *et al*. 2015) with sample type as a factor and bird ID as a random intercept to account for the dependence of observations between duplicates and samples. Repeatability of traits (*R*) for each sample type was expressed as the intraclass correlation coefficient with 95% confidence intervals based on 1000 bootstrap iterations using the rptR package (Stoffel, Nakagawa & Schielzeth 2017). We used lmer models to address the effect of sample volume on respiration rates and FCRs (i.e., Experiment 2) using the same general model structure. Predicted values and their standard errors were calculated using the emmeans package (Lenth 2019) and used the pairs function in this package to perform post hoc tests for Experiment 2 data. The effects of storage time (i.e., Experiment 3) were assessed using paired t-tests between the 0 h and 24 h samples (t.test function in R base).

## RESULTS

A representative trace of oxygen consumption in a whole blood experiment is provided in the Electronic Supplementary Materials (Fig. S1 in ESM1).

### Effects of sample type – blood cells vs. whole blood

ROUTINE respiration did not differ between isolated blood cell and whole blood samples (Table 1, Fig. 1A). However, OXPHOS and ETS were significantly higher when measured on whole blood samples compared to on blood cell isolates (1.2 and 1.3-fold, respectively) (Figs 1B-C, Table 1). In contrast, LEAK was 30% lower in the whole blood samples (Fig. 1D, Table 1). Consequently, the ER control efficiency (which measures proportionally how much respiration can be increased from ROUTINE) was significantly higher in the whole blood samples (0.511, compared to 0.365 in isolated cells; Fig. 2A, Table 1). Whole blood samples also had significantly higher phosphorylating efficiency and higher coupling efficiency during maximal FCCP-stimulated respiration (i.e., R-L control and E-L coupling efficiencies were higher) (Fig. 2B-C, Table 1).

**Table 1.**
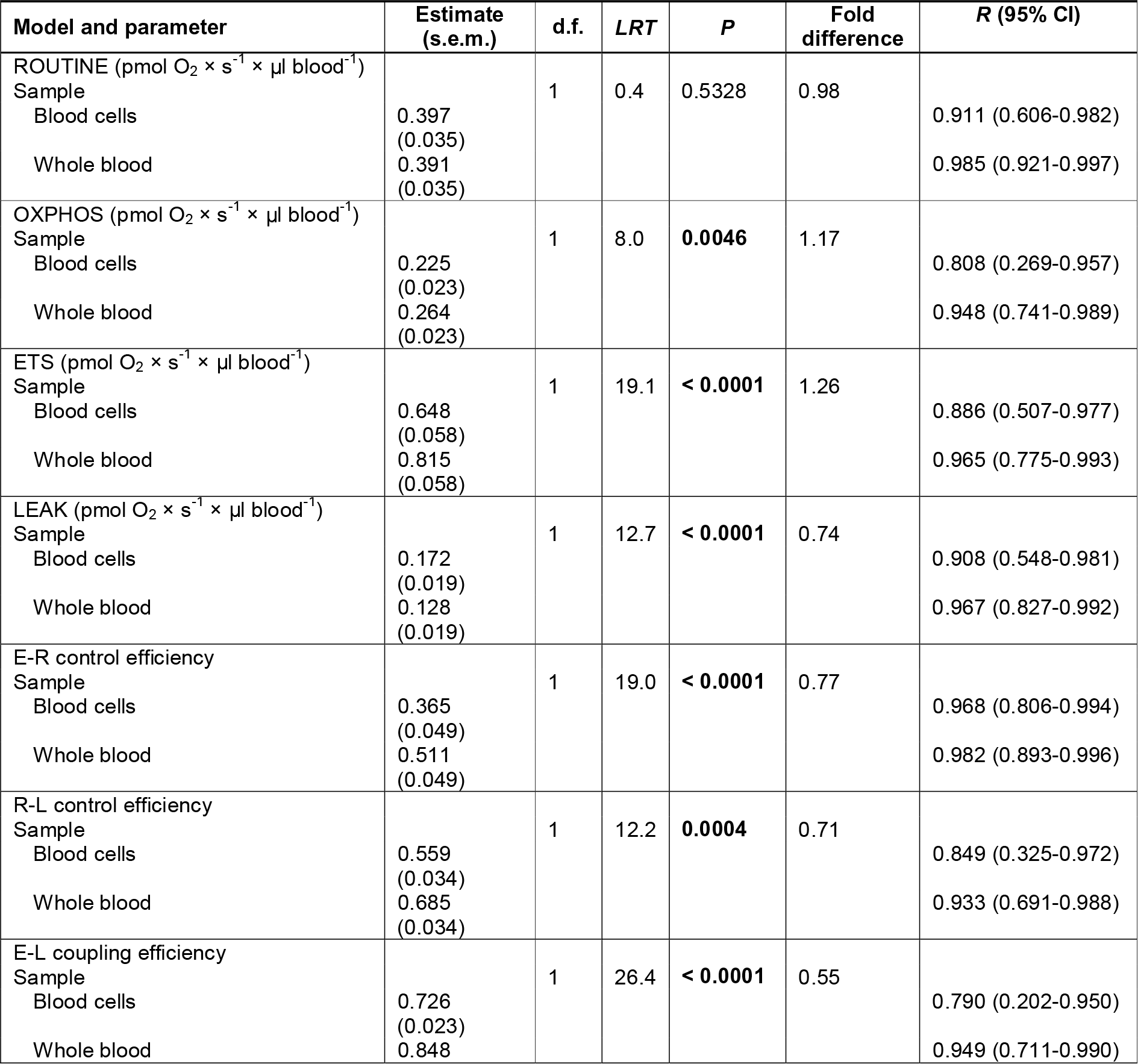
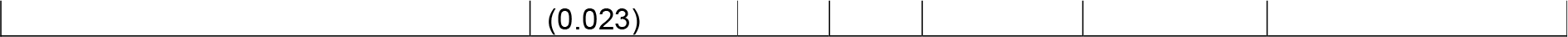
Comparison of mitochondrial respiration metrics in whole blood and in isolated blood cells from the same blood volume. The table shows model estimates, test statistics, degrees of freedom and resultant *P*-values, when assaying mitochondrial respiration in zebra finches using either 100 μl whole blood or all cells from 100 μl blood. All samples were run in duplicate. *R* refers to intraclass repeatability of these samples and is presented with 95% confidence bands inferred from 1000 bootstrap iterations. Significant (i.e., *P* < 0.05) effects appear in bold font. Abbreviations: CI = Confidence Interval; d.f. = Degrees of Freedom; *LRT* = Likelihood Ratio Test statistic; *R* = Intraclass Repeatability; s.e.m. = Standard Error of Mean.

| Model and parameter | Estimate (s.e.m.) | d.f. | *LRT* | *P* | Fold difference | *R* (95% CI) |
| --- | --- | --- | --- | --- | --- | --- |
| ROUTINE (pmol O<sub>2</sub> × s<sup>-1</sup> × μl blood<sup>-1</sup>) | | | | | | |
| Sample |  | 1 | 0.4 | 0.5328 | 0.98 |  |
| Blood cells | 0.397 (0.035) |  |  |  |  | 0.911 (0.606-0.982) |
| Whole blood | 0.391 (0.035) |  |  |  |  | 0.985 (0.921-0.997) |
| OXPHOS (pmol O<sub>2</sub> × s<sup>-1</sup> × μl blood<sup>-1</sup>) | | | | | | |
| Sample |  | 1 | 8.0 | **0.0046** | 1.17 |  |
| Blood cells | 0.225 (0.023) |  |  |  |  | 0.808 (0.269-0.957) |
| Whole blood | 0.264 (0.023) |  |  |  |  | 0.948 (0.741-0.989) |
| ETS (pmol O<sub>2</sub> × s<sup>-1</sup> × μl blood<sup>-1</sup>) | | | | | | |
| Sample |  | 1 | 19.1 | **< 0.0001** | 1.26 |  |
| Blood cells | 0.648 (0.058) |  |  |  |  | 0.886 (0.507-0.977) |
| Whole blood | 0.815 (0.058) |  |  |  |  | 0.965 (0.775-0.993) |
| LEAK (pmol O<sub>2</sub> × s<sup>-1</sup> × μl blood<sup>-1</sup>) | | | | | | |
| Sample |  | 1 | 12.7 | **< 0.0001** | 0.74 |  |
| Blood cells | 0.172 (0.019) |  |  |  |  | 0.908 (0.548-0.981) |
| Whole blood | 0.128 (0.019) |  |  |  |  | 0.967 (0.827-0.992) |
| E-R control efficiency |  |  |  |  |  |  |
| Sample |  | 1 | 19.0 | **< 0.0001** | 0.77 |  |
| Blood cells | 0.365 (0.049) |  |  |  |  | 0.968 (0.806-0.994) |
| Whole blood | 0.511 (0.049) |  |  |  |  | 0.982 (0.893-0.996) |
| R-L control efficiency |  |  |  |  |  |  |
| Sample |  | 1 | 12.2 | **0.0004** | 0.71 |  |
| Blood cells | 0.559 (0.034) |  |  |  |  | 0.849 (0.325-0.972) |
| Whole blood | 0.685 (0.034) |  |  |  |  | 0.933 (0.691-0.988) |
| E-L coupling efficiency |  |  |  |  |  |  |
| Sample |  | 1 | 26.4 | **< 0.0001** | 0.55 |  |
| Blood cells | 0.726 (0.023) |  |  |  |  | 0.790 (0.202-0.950) |
| Whole blood | 0.848 |  |  |  |  | 0.949 (0.711-0.990) |

**Figure 1.**
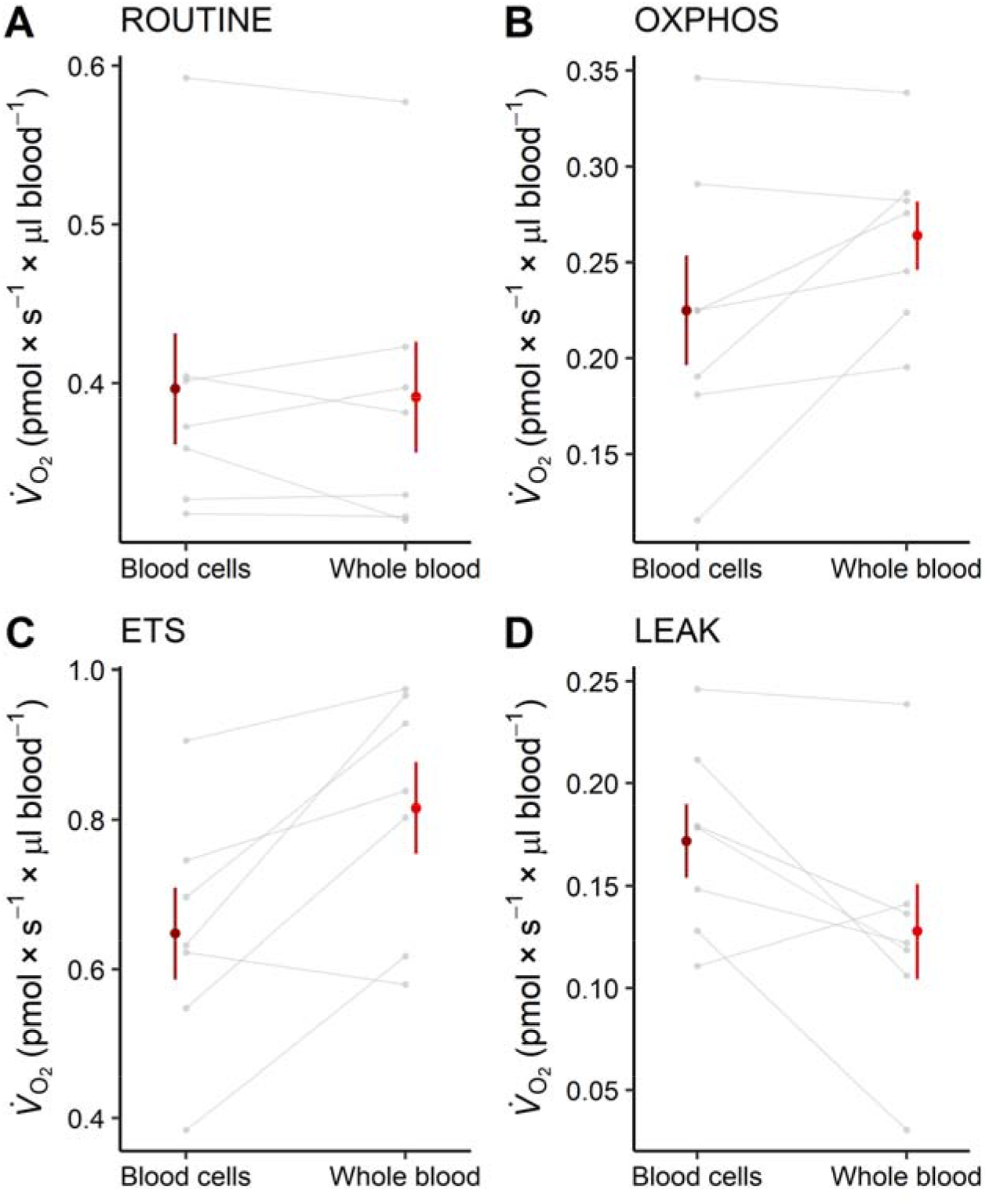
Difference in mitochondrial respiration traits as assayed in either isolated blood cells or whole blood from zebra finches. A) baseline (“ROUTINE”) oxygen consumption in the endogenous cellular state; B) oxygen consumption devoted to oxidative phosphorylation alone, where ATP is produced (“OXPHOS); C) maximum working capacity of the electron transport system when uncoupled from ATP production (“ETS”); D) the part of ROUTINE used to offset the leak of protons across the inner mitochondrial membrane (“LEAK). Coloured plotting symbols show raw data means ± 1 standard error. Gray points and lines show individual responses. Significances are reported in Table 1.

**Figure 2.**
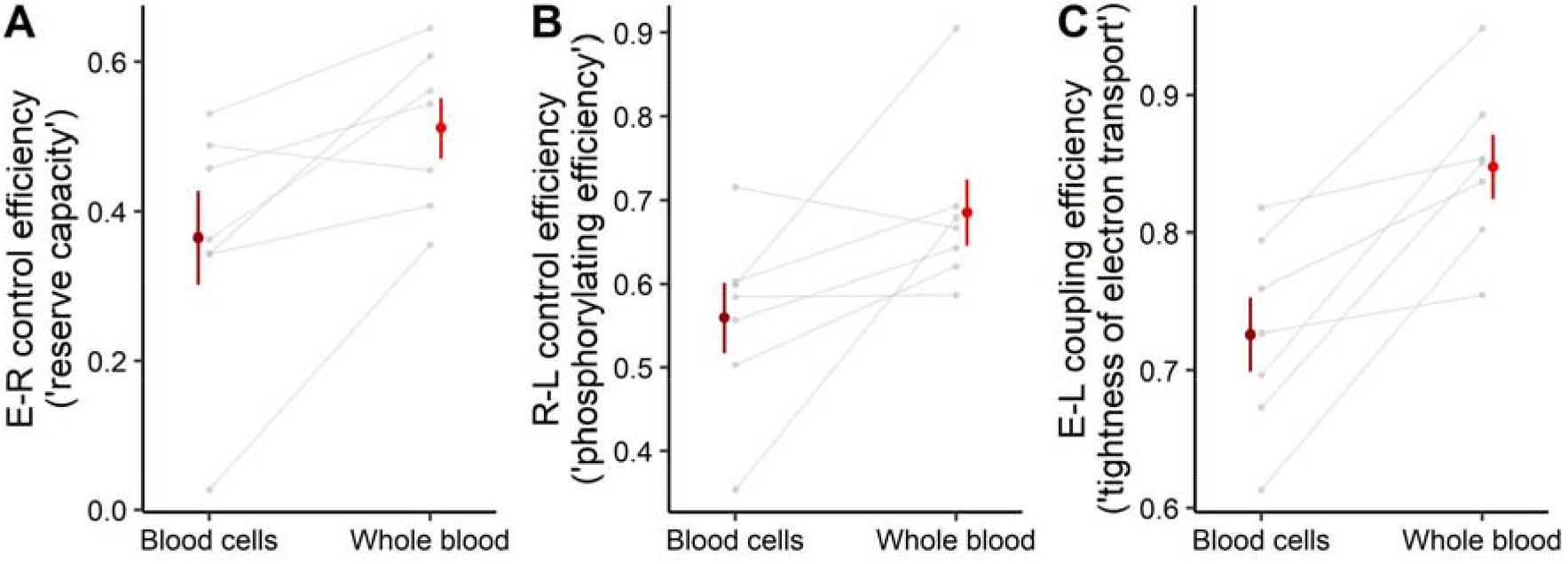
Flux Control Ratios for mitochondrial respiration traits measured in isolated blood cells or whole blood. A) The proportion of maximum working capacity (ETS) remaining during endogenous respiration (ROUTINE); B) Efficiency of ATP-producing respiration (OXPHOS) under endogenous cellular conditions; C) Coupling efficiency of electron transport in a FCCP-stimulated cellular state. Coloured plotting symbols show raw data means ± 1 standard error. Gray points and lines show individual responses. Details on calculation of FCRs are provided in the main text, and significances appear in Table 1.

Respiration traits measured on both whole blood and isolated blood cells were significantly and highly reproducible (blood cells: *R*_min_ = 0.790, *R*_maz_ = 0.968; whole blood: *R*_min_ = 0.849, *R*_max_ = 0.985). However, whole blood samples were more reproducible, and the calculated repeatabilities had markedly narrower confidence intervals, for all respiration traits and FCRs investigated (Table 1).

### Effects of blood volume

There were non-linear effects on blood sample volume on respiration traits and FCRs, with the lowest and highest sample volumes showing the most pronounced deviations. Accordingly, ROUTINE was significantly higher (by a factor 1.5 to 1.6) for all volumes 10-50 μl compared to the 100 μl sample (Fig. 3A, Table 2). OXPHOS was the highest, and similar for 25 and 50 μl, and the lowest for 10 and 100 μl (which did not differ) (Fig. 3A, Table 2). OXPHOS measured using 50 μl was 1.3-fold higher than the measurement using 10 μl, though this difference was not statistically significant. ETS was the highest on 50 μl samples (up to 1.4-fold), and significantly different from all volumes but 25 μl (despite a 1.2-fold difference; Table 2). ETS was the lowest on 100 μl samples, though this volume differed statistically only from 50 μl (Table 2). LEAK was significantly higher on 10 μl compared to all other volumes, and the lowest on 100 μl (though this volume was not statistically different from 50 μl) (Table 2).

**Table 2.** Effects of sample volume when measuring mitochondrial respiration in zebra finch whole blood. The table shows model estimates (± standard error; s.e.m.), test statistics, degrees of freedom and corresponding *P*-values when comparing the effect of sample volume when measuring mitochondrial respiration in zebra finch whole blood. Significant (i.e., *P* < 0.05) effects appear in bold font. Different letters with brackets denote statistically significant pairwise differences. Abbreviations: d.f. = Degrees of Freedom; *LRT* = Likelihood Ratio Test statistic; s.e.m. = Standard Error of Mean. \**P*_10–25 μl_ = 0.053.

| Model and parameter | Estimate<br>(s.e.m.) | d.f. | *LRT* | *P* |
| --- | --- | --- | --- | --- |
| ROUTINE (pmol O<sub>2</sub> × s<sup>-1</sup> × μl blood<sup>-1</sup>)<br>Blood volume<br>10 μl [A]<br>25 μl [A]<br>50 μl [A]<br>100 μl [B] | 0.669<br>(0.037)<br>0.656<br>(0.034)<br>0.607<br>(0.034)<br>0.411<br>(0.034) | 1 | 27.5 | **< 0.0001** |
| OXPHOS (pmol O<sub>2</sub> × s<sup>-1</sup> × μl blood<sup>-1</sup>)<br>Blood volume<br>10 μl [AB]<br>25 μl [C]*<br>50 μl [BC]<br>100 μl [A] | 0.328<br>(0.029)<br>0.437<br>(0.026)<br>0.416<br>(0.026)<br>0.259<br>(0.026) | 1 | 22.2 | **< 0.0001** |
| ETS (pmol O<sub>2</sub> × s<sup>-1</sup> × μl blood<sup>-1</sup>)<br>Blood volume<br>10 μl [A]<br>25 μl [AB]<br>50 μl [B]<br>100 μl [A] | 0.784<br>(0.060)<br>0.835<br>(0.055)<br>1.005<br>(0.055)<br>0.715<br>(0.055) | 1 | 13.9 | **0.0031** |
| LEAK (pmol O<sub>2</sub> × s<sup>-1</sup> × μl blood<sup>-1</sup>)<br>Blood volume<br>10 μl [A]<br>25 μl [B]<br>50 μl [BC]<br>100 μl [C] | 0.345<br>(0.018)<br>0.219<br>(0.017)<br>0.191<br>(0.017)<br>0.152<br>(0.017) | 1 | 39.9 | **< 0.0001** |
| E-R control efficiency<br>Blood volume<br>10 μl [A]<br>25 μl [A]<br>50 μl [B]<br>100 μl [B] | 0.159<br>(0.051)<br>0.191<br>(0.049)<br>0.389<br>(0.049)<br>0.421<br>(0.049) | 1 | 29.6 | **< 0.0001** |
| R-L control efficiency<br>Blood volume<br>10 µl [A] | 0.486<br>(0.025) | 1 | 28.5 | **< 0.0001** |
| 25 µl [B] | 0.664<br>(0.022) |  |  |  |
| 50 µl [B] | 0.685<br>(0.022) |  |  |  |
| 100 µl [B] | 0.625<br>(0.022) |  |  |  |
| E-L coupling efficiency<br>Blood volume<br>10 µl [A] | 0.566<br>(0.023) | 1 | 49.7 | **< 0.0001** |
| 25 µl [B] | 0.726<br>(0.022) |  |  |  |
| 50 µl [C] | 0.803<br>(0.022) |  |  |  |
| 100 µl [C] | 0.785<br>(0.022) |  |  |  |

**Figure 3.**
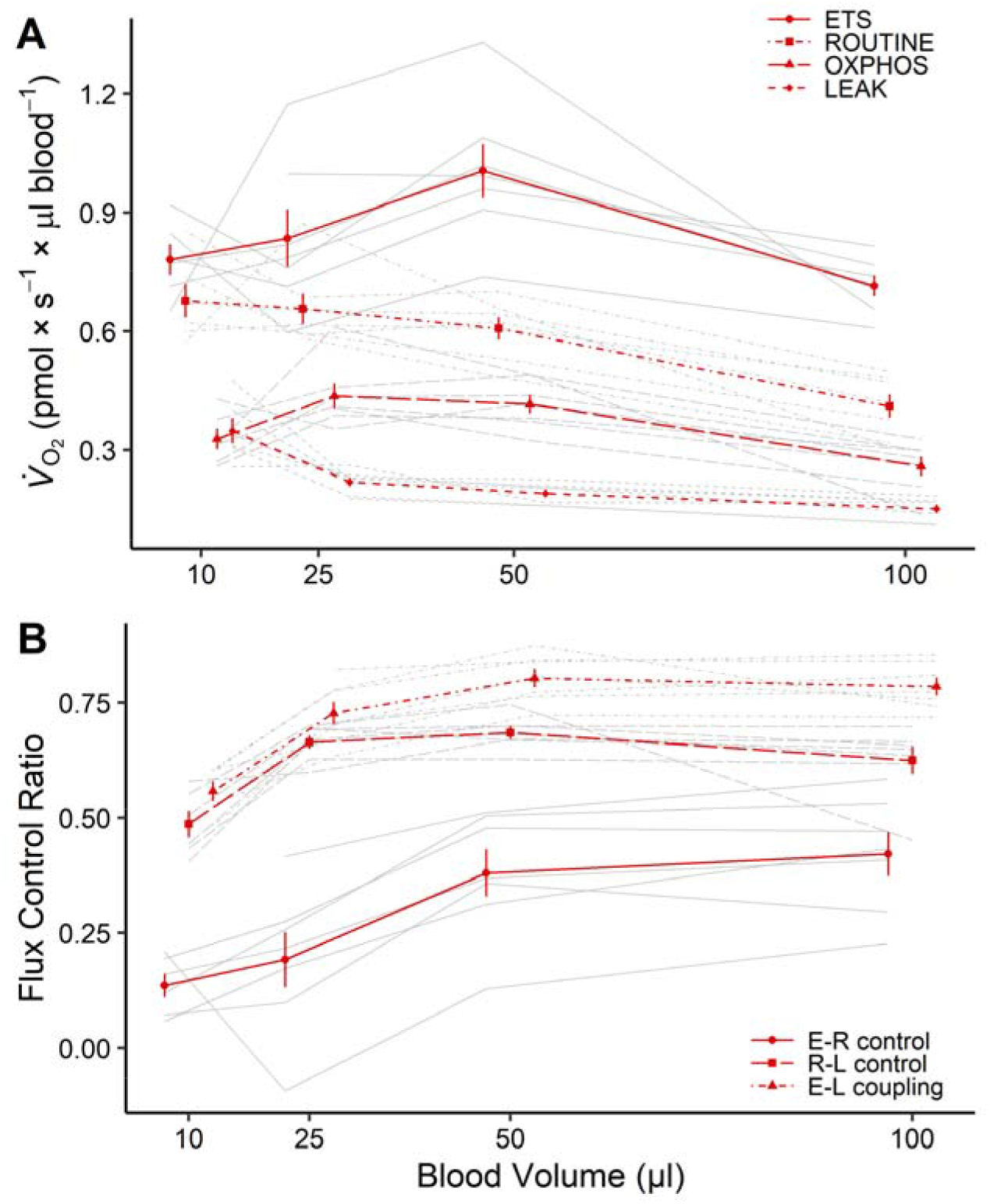
Effects of sample volume when assaying mitochondrial function in zebra finch whole blood. The panels show mitochondrial respiration traits (A) and Flux Control Ratios (FCR) (B) in relation to blood sample volume. Derivation of traits and FCRs are detailed in the main text and in the legend to Fig. 2. Coloured plotting symbols show raw data means ± 1 standard error, and coloured lines show the averaged response. Gray lines show individual responses. Significances are reported in Table 2.

FCRs, which provide rate-independent insight into mitochondrial function, were also volume-specific. Specifically, E-R control efficiency was similar in 50 and 100 μl and significantly higher (reflecting more surplus capacity) compared to the two other volumes (Fig. 3B). R-L control efficiency (i.e., the efficiency with which O_2_ consumption is channelled towards oxidative phosphorylation) was the lowest in 10 μl (reflecting higher LEAK) compared to the other volumes (Table 2, Fig. 3B). E-L coupling efficiency was also the lowest in 10 μl followed by 25 μl, but higher in both 50 and 100 μl (which were not statistically different) (Table 2, Fig. 3B).

### Effects of storage

Endogenous (ROUTINE), phosphorylating (OXPHOS) and leak (LEAK) respiration did not change significantly with 24 h storage at 5°C (all *P* > 0.2) (Fig. 4A, Table 3). However maximal respiration capacity (ETS) was 16% lower at 24 h compared to at 0 h. As a result, E-R control efficiency and E-L coupling efficiency was significantly lower at 24 h (Fig. 4B, Table 3). However, there was no change in R-L control efficiency, in keeping with stable ROUTINE and LEAK over 24 h storage. By 72 h, ROUTINE, OXPHOS and ETS had decreased somewhat from the 24 h level and LEAK had increased, with corresponding changes to FCRs (Fig. S2 in ESM2). These changes became more exaggerated with 96 h storage (Fig. S2).

**Table 3.** Effects of storage time on mitochondrial respiration metrics in zebra finch whole blood. The table shows mean (± standard error; s.e.m.) trait values at 0 h and after 24 h storage at 5°C, as well as test statistics, degrees of freedom and corresponding *P*-values for paired *t*-tests of the effects of storage time on respiration traits. Significant (i.e., *P* < 0.05) effects appear in bold font. Abbreviations: d.f. = Degrees of Freedom; s.e.m. = Standard Error of Mean; *t* = *t-*test statistic.

| Model and parameter | Mean (s.e.m.) | d.f. | *t* | *P* |
| --- | --- | --- | --- | --- |
| ROUTINE (pmol O<sub>2</sub> × s<sup>-1</sup> × µl blood<sup>-1</sup>)<br>Sample<br>0 h<br>24 h | 0.673<br>(0.047)<br>0.731<br>(0.047) | 5 | -1.40 | 0.2214 |
| OXPHOS (pmol O<sub>2</sub> × s<sup>-1</sup> × µl blood<sup>-1</sup>)<br>Sample<br>0 h<br>24 h | 0.498<br>(0.046)<br>0.542<br>(0.035) | 5 | -1.47 | 0.2015 |
| ETS (pmol O<sub>2</sub> × s<sup>-1</sup> × µl blood<sup>-1</sup>)<br>Sample<br>0 h<br>24 h | 1.169<br>(0.088)<br>1.011<br>(0.111) | 5 | 3.23 | **0.0233** |
| LEAK (pmol O<sub>2</sub> × s<sup>-1</sup> × µl blood<sup>-1</sup>)<br>Sample<br>0 h<br>24 h | 0.175<br>(0.008)<br>0.188<br>(0.019) | 5 | -0.94 | 0.3905 |
| E-R control efficiency<br>Sample<br>0 h<br>24 h | 0.404<br>(0.066)<br>0.255<br>(0.049) | 5 | 4.16 | **0.0089** |
| R-L control efficiency<br>Sample<br>0 h<br>24 h | 0.736<br>(0.019)<br>0.744<br>(0.016) | 5 | -0.74 | 0.4904 |
| E-L coupling efficiency<br>Sample<br>0 h<br>24 h | 0.845<br>(0.017)<br>0.809<br>(0.016) | 5 | 3.29 | **0.0217** |

**Figure 4.**
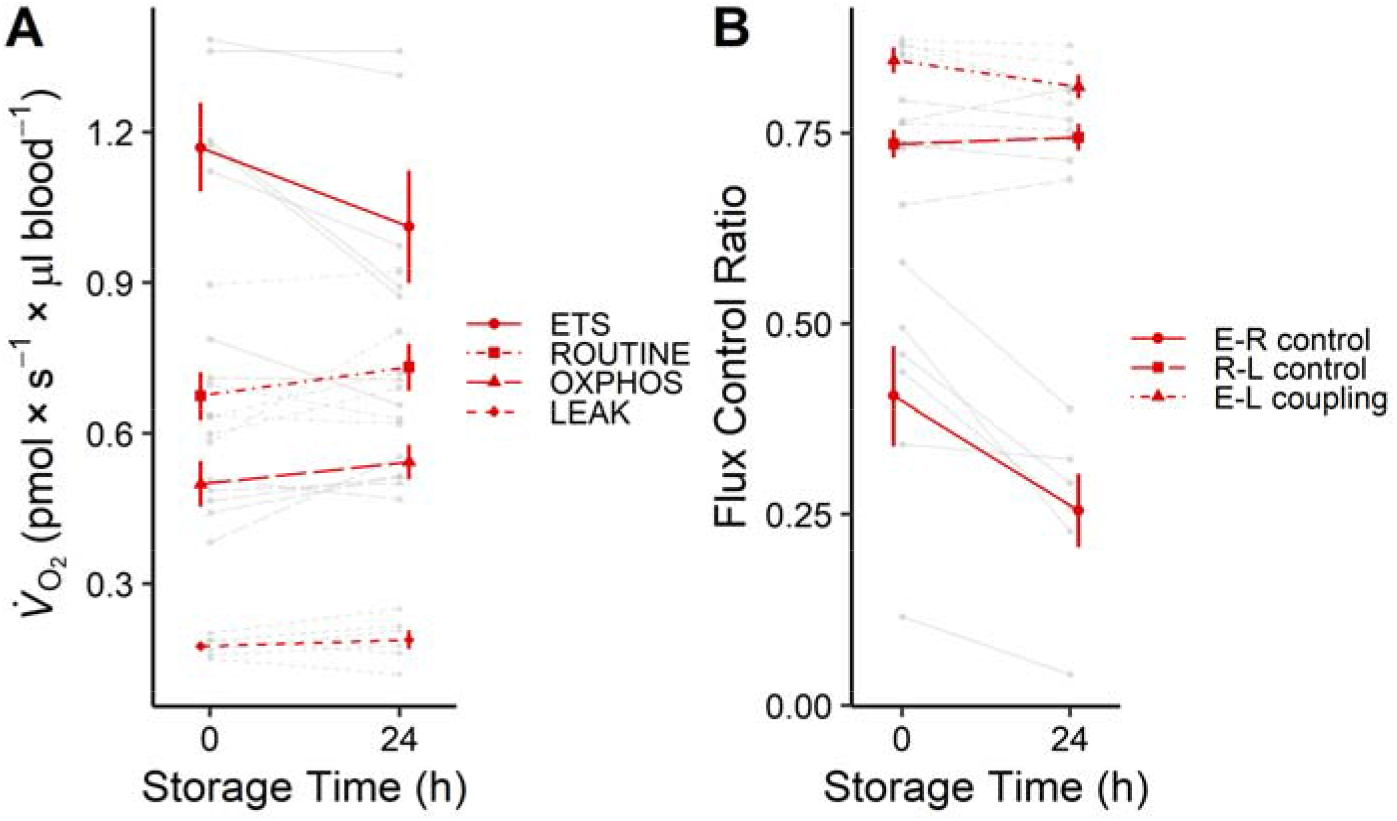
Effects of storage time when assaying mitochondrial function in zebra finch whole blood. The panels show mitochondrial respiration traits (A) and Flux Control Ratios (FCR) (B) in relation to blood sample volume. All individuals were measured at 0 h and after storage without agitation for 24 h at 5°C. Derivation of traits and FCRs are detailed in the main text and in the legend to Fig. 2. Coloured plotting symbols show raw data means ± 1 standard error, and coloured lines show the averaged response. Gray lines show individual responses. Significances are reported in Table 3.

## DISCUSSION

In this study, we asked whether measurement on whole blood could provide a feasible and potentially more biologically contextualised approach to studying mitochondrial respiration in intact avian blood cells. While endogenous respiration did not differ between whole blood samples and isolated cells produced from the same blood volume, we found that both phosphorylating (i.e., OXPHOS; Fig. 1B) and maximal (i.e., ETS; Fig. 1C) respiration was significantly higher in whole blood, and that mitochondria in isolated blood cells displayed significantly more LEAK (i.e., they were more uncoupled) than whole blood (Fig. 1D). This translated to significant differences in flux control ratios (FCRs), which indicated that the isolated blood cells had lower surplus respiratory capacity (Fig. 2A), lower phosphorylating efficiency (Fig. 2B) and reduced coupling efficiency when stimulated to work at maximum (Fig. 2C). Moreover, while our repeatability estimates for isolated erythrocytes were consistently high and mostly within the range for the same traits reported elsewhere (Stier et al. 2017), the whole blood samples were consistently more reproducible (all *R* > 0.93) and less variable for all traits considered (Table 1). Thus, in our study whole blood measurement was more precise, and biological inference differed in several key traits depending on whether measurements were performed on the intact tissue or on a derived sample. We believe that this could be caused by physical damage or reduced integrity of mitochondria in the isolated blood cells caused by more handling, because lower ETS, higher LEAK and associated changes to FCRs in repeated samples of the same subject are all indicative of declining mitochondrial health/viability (e.g. Sjövall *et al*. 2013a) (see also Fig. 4). Studies on mammalian erythrocytes show centrifugation alone can cause time-dependent structural and physiological damage to red blood cells (such as increased free and corpuscular haemoglobin and increased ATP release) at spinning speeds considerably lower than those used here (< 1500 *g*) (Kiss *et al*. 2016; Urbina *et al*. 2016; Wiegmann *et al*. 2017; Mancuso, Jayaraman & Ristenpart 2018). We are not suggesting that our whole blood samples were in a ‘pristine’ condition, because drawing blood into a syringe, and subsequent pipetting of a blood sample, may also impact the physiological and physical state of the red blood cells (Wiegmann *et al*. 2017). However, our study indicates that whole blood measurement may salvage more of the physiological integrity of the avian blood cell.

We found non-linearity of volume-adjusted respiration rates, manifested as inhibited or enhanced respiration on the highest and lowest blood volumes (Fig. 3A). On average, optimal trait combinations for coupled and uncoupled respiration were achieved using 50 μl samples, but these runs were most often comparable to those on 25 μl. FCR for phosphorylating and coupling efficiencies were similar in the range of 25 to 100 μl, whereas 10 μl samples were less well coupled. Thus, working in a range of 25 to 50 μl whole blood provided representative results for respiration traits, whereas a sample volume range of 25 to 100 μl was acceptable if FCRs only were considered. Our findings contrast previous studies on pig liver homogenates where complete linearity of phosphorylating respiration was demonstrated across a 3-fold range in sample volume (Kuznetsov *et al*. 2002). However, studies on isolated human platelets show reduced uncoupled respiration on small sample volumes (Sjövall *et al*. 2013a). This could be caused e.g., by reduced accuracy on low sample concentrations acting in combination with enhanced sensitivity/exposure to inhibitors. Measurements on pig skeletal muscle mitochondria also suggest suppressed respiration when samples were prepared from smaller biopsies (Isner-Horobeti *et al*. 2014), though this probably reflects heterogeneity in sample composition that was not accounted for by weight alone. It is less clear why we observed inhibition of respiration (but not FCRs) on large blood sample volumes. We do not believe that this was caused by the presence of plasma in the sample for at least two reasons: 1) both intact and open blood cells can be measured using plasma as medium with inhibition of respiration (Sjövall *et al*. 2010; Grip *et al*. 2016); 2) ROUTINE was identical in whole blood and isolated blood cells (Fig. 1A), which is not expected if some whole blood constituents act suppressively. The explanation for these observations should be addressed in future work, including measurement of mitochondrial content in differently sized samples, and using both intact and open-cell protocols. Meanwhile, we recommend experimenters to use relatively constant sample volumes to the furthest extent possible.

The whole blood samples were robust to cold storage. After 24 h at 5°C without agitation, all respiration traits but ETS (which decreased by 14%) remained unaltered. This corroborates previous work on both human and avian blood cells (Sjövall *et al*. 2013a; Stier *et al*. 2017) and pig liver homogenates (Kuznetsov *et al*. 2002). However, neither Sjövall et al. (2013) or Stier et al. (2017) found any reduction in ETS over 24 h. These authors measured the cells 3-5 h after blood collection, whereas our first measurement was performed within 45 min of bleeding the birds. It is tempting to speculate that most changes to ETS occur within the first hour of collection. Sjövall (2013) found that respiration of permeabilised human platelets remained largely unaltered up to 48 h when the blood was stored at room temperature. This does not seem to be the case in birds: while ROUTINE, OXPHOS and LEAK remained relatively stable when the blood was stored 2-2.5 h at room temperature under constant motion, ETS dropped by 30 to 90% over the same period (data not shown).

## CONCLUSIONS

We found that whole blood measurement provides more rapid, more biologically contextualised, and more precise and reproducible assessment of mitochondrial respiration in intact avian hematocytes compared to measurement on isolated cells. We also give data-based recommendations on sample volume ranges, show that 24 h cold-storage in the blood collection tubes is possible at acceptable changes to respiration traits and that whole blood can be permeabilised. Changes to respiration parameters of the intact cells were indicative of handling-induced damage. However, it is possible that less extensive handling protocols (see e.g. Stier *et al*. 2019) will provide more comparable results. Even so, our study suggests that centrifugation and removal of plasma are unnecessary complications that add processing time and reduce biological context. Thus, we advocate use of whole blood whenever possible. When measurements on isolated cells are necessary, such as when low sample volume is not sufficient for complementary analyses of plasma metabolites, care should be taken to keep mechanical stress to a minimum. We also recommend use of relatively constant blood volumes to minimize variation within an experiment, and suggest samples are run on the day of collection.

## Supporting information

Electronic Supplementary Material 1

Electronic Supplementary Material 2

## ACKNOWLEDGEMENTS

The authors thank Elsie Ye Xiong and Michael Tobler for providing zebra finches for use in the study. Camilla Björklöv and Agnieszka Czopek excellently assisted with care for the experimental animals. Comments from Elisa Thoral improved a previous version of the manuscript.

## ETHICS

All procedures were approved by the Malmö/Lund Animal Ethics Committee, acting under authority of the Swedish Board of Agriculture (permit no. 9246-19, 16977-19).

## DATA AVAILABILITY

Data will be deposited in Figshare upon acceptance of the manuscript.

