## Supplementary material for "A whole blood approach improves speed and accuracy when measuring mitochondrial respiration in intact avian hematocytes": Electronic Supplementary Material 1

### Developing an open cell protocol for avian whole blood

Intact cell protocols allow insights into mitochondrial respiration under endogenous cellular conditions, but by necessity does not permit in-depth understanding of mitochondrial complex function, such as respiration on varying amounts of substrates or responses to different substrate combinations. Thus, we explored the possibility of permeabilising blood cells in whole blood samples. Blood was drawn and processed as above (*n* = 3; 2 females, 1 male). We ran 3 samples of 25 µl per bird simultaneously, having previously established that this sample volume yielded an optimal balance between phenotypic responses and $O_2$ saturation during the open cell run (data not shown). Once we had collected 2-3 min of stable ROUTINE respiration, we added 0, 5, or 10 µl 5 mg ml$^{-1}$ of the detergent digitonin (i.e., 0, 0.2 or 0.4 µg per µl blood) to permeabilise blood cell membranes. Using different concentrations of the permeabilising agent allowed us to estimate which level of permeabilization that yielded the biologically most relevant results (details below). Four to 5 minutes later, we added 5 mM malate and 5 mM pyruvate to fuel Complex I. When respiration on malate and pyruvate had remained stable for 2-3 min, we added 1 mM ADP to trigger phosphorylating respiration via Complex I. This allowed us to calculate Complex I activity without (henceforth 'State 2') and with (henceforth '$OXPHOS_{CI}$') concomitant ATP production. The additional Complex I-linked substrate glutamate (5 mM) never increased respiration above $OXPHOS_{CI}$ and so these data are not presented below. We then added 10 mM succinate, which is used by Complex II, to elicit maximum phosphorylating respiration on excess substrate (henceforth '$OXPHOS_{CI+II}$'). We subsequently induced LEAK state by inhibiting ATP synthase using 1 µg ml$^{-1}$ oligomycin (henceforth '$LEAK_{Omy}$'), followed by uncoupling of the electron transport system by titration of 0.5 µl aliquots of 1 mM FCCP until maximum (henceforth '$ETS_{CI+II}$') (final concentration: 1-2 µM). Afterwards, we added 2 μM rotenone to measure the contribution of Complex II to ETS (henceforth ‘$ETS_{CII}$’). Finally, we added 1 μg $ml^{-1}$ antimycin A to gauge non-mitochondrial respiration.

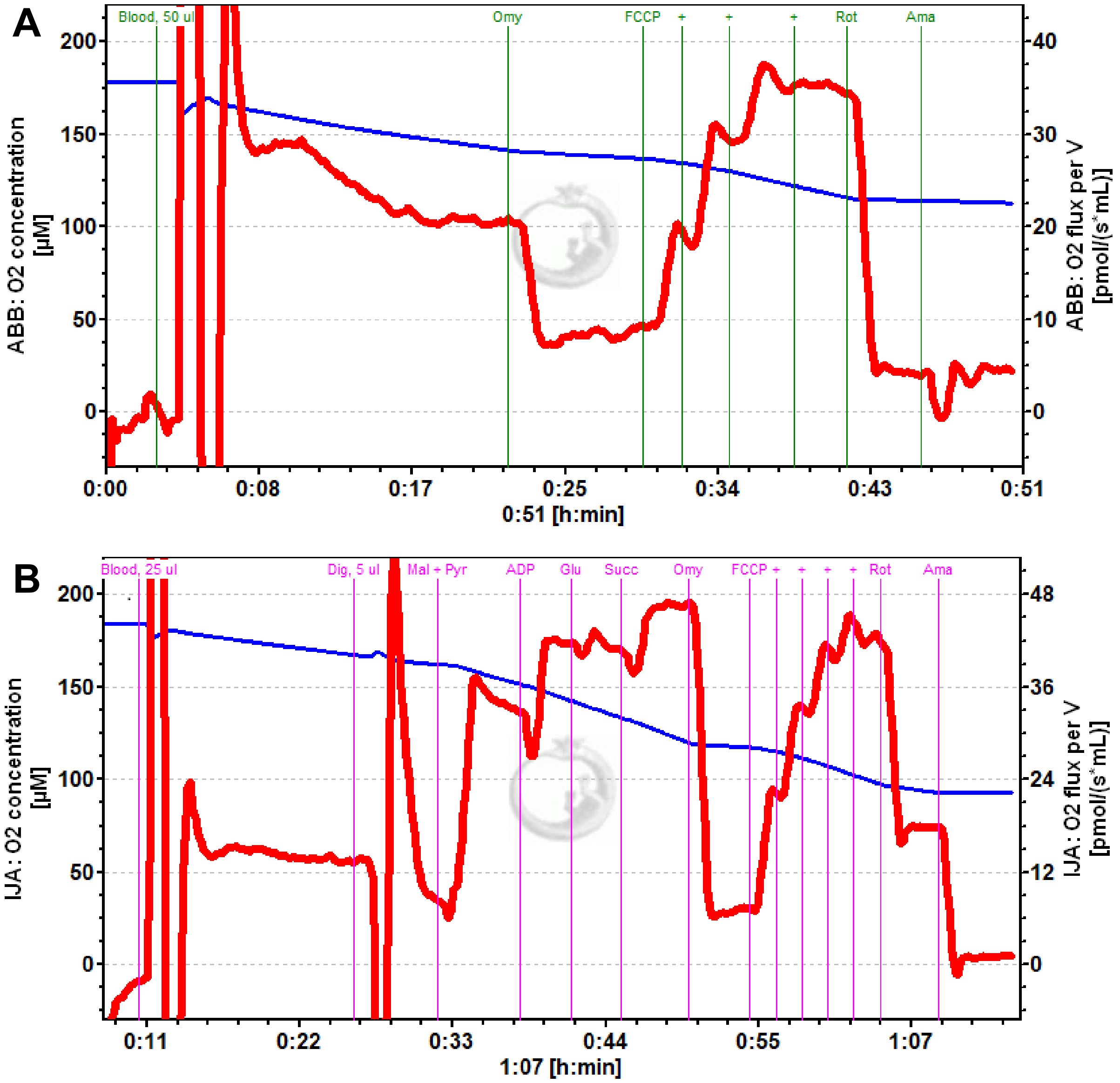


**Figure S1. Representative traces of intact and open cell protocols when measuring mitochondrial respiration in bird whole blood.** A) Intact cell measurement of 50 µl whole blood from an individual zebra finch (*Taeniopygia guttata* Vieillot) female (individual 5716C). B) Open cell measurement of 25 µl whole blood permeabilised with 5 µl 5 mg $ml^{-1}$ digitonin from an individual zebra finch female (individual 5416C). The red curve shows oxygen flux in the respirometry chamber, and the blue curve shows oxygen concentration during the experiment. Birds and blood samples were handled as described in the main text. Concentrations of substrates and inhibitors are detailed in the main text. Abbreviations: Dig = Digitonin; Mal = Malate; Pyr = Pyruvate; ADP = Adenosine diphosphate; Glu = Glutamate; Succ = Succinate; Omy = Oligomycin; FCCP = Carbonyl cyanide 4-(trifluoromethoxy)phenylhydrazone; + = Subsequent FCCP titrations; Rot = Rotenone; Ama = Antimycin A.

Digitonin readily permeabilised the blood cells in whole blood samples (Fig. S1). The responses to 0.2 and 0.4 µg digitonin per µl blood yielded comparable estimates of phosphorylating (i.e., $OXPHOS_{CI}$, $OXPHOS_{CI+II)}$, uncoupled (i.e., $ETS_{C1}$, $ETS_{C1+2}$) and $LEAK_{Omy}$ respiration (Fig. S2). On average, all respiration traits were markedly higher in the presence of digitonin compared to when no digitonin was added. However, maximal uncoupled respiration (i.e., $ETS_{C1+2}$) was broadly similar in the experiments with and without digitonin (Fig. S2).

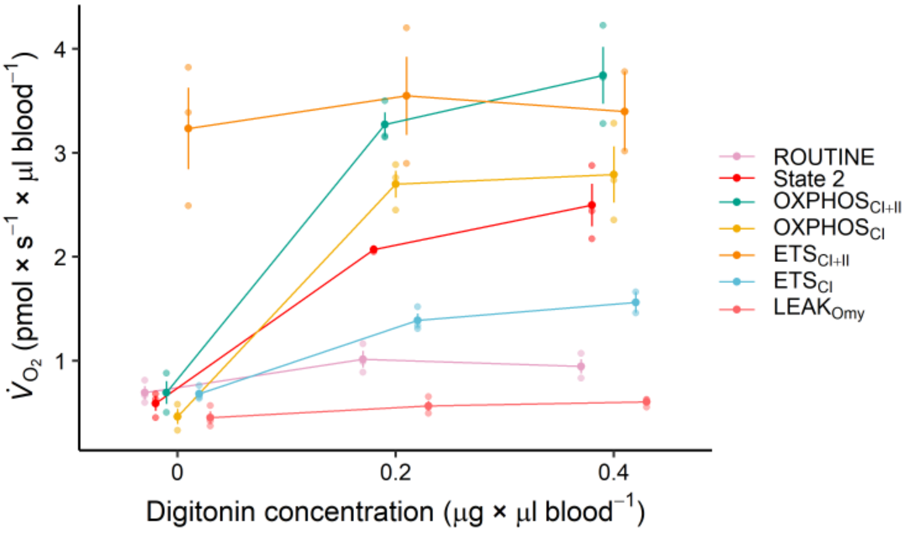


**Figure S2. Effect of digitonin concentration on mitochondrial respiration traits measured in zebra finch whole blood samples.** The figure shows how variation in concentration of the permeabilising detergent digitonin (5 mg ml$^{-1}$) affected mitochondrial respiration traits in hematocytes from 25 µl whole blood samples collected from captive zebra finches. Calculation of respiratory states is explained in the main text. Solid plotting symbols lines show raw data means ± 1 standard error, and coloured lines show the averaged response. The smaller, semi-transparent plotting symbols show raw data.

Our pilot study suggests that hematocytes contained in whole blood can be permeabilised to gain further insights into mitochondrial function, comparable to the case in isolated blood cells (Stier et al. 2019). We assumed that the most biologically relevant level of permeabilization would be characterised by stronger relative responses to ADP (i.e., an increase in $OXPHOS_{CI}$ / State 2) and greater similarity between maximal uncoupled and phosphorylating respiration (i.e., an increase in $OXPHOS_{CI+II}$ / $ETS_{CI+II}$). Based on this $OXPHOS_{CI}$ capacity was the highest for 0.2 µg digitonin per µl blood. However, dose responses need to be determined in dedicated studies. Finally, it is noteworthy that addition of substrates to intact cells yielded nearly identical values for $OXPHOS_{CI+II}$ compared to the open cells, driven mostly by Complex II activity). This semi-permeability to substrates in avian blood cells could potentially be exploited in various intact cell applications both *in vitro* and *in vivo*.
