## Supplementary material for "A whole blood approach improves speed and accuracy when measuring mitochondrial respiration in intact avian hematocytes": Electronic Supplementary Material 2

### Effects of blood storage beyond 24 hours

The blood samples from two females were opportunistically sampled at each of 72 h and 96 h after collection using the methods outlined in the main text. This revealed a slight reduction in ROUTINE, OXPHOS and ETS, and an increase in LEAK, by 72 h. Flux control ratios changed accordingly. These changes became more exaggerated with 96 h storage (Fig. S2).

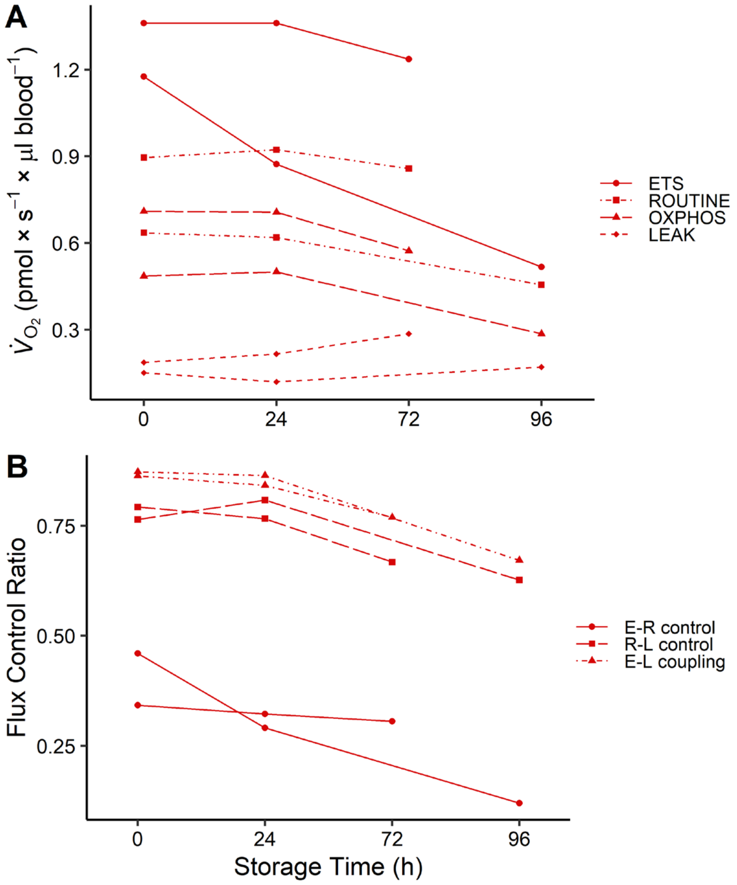


**Figure S2. Effects of cold storage beyond 24 hours when assaying mitochondrial respiration in bird whole blood.** Data were collected from two female zebra finches (*Taeniopygia guttata* Vieillot) that were opportunistically re-measured at 72 h and 96 h after blood sample collection. Panel A) shows mitochondrial respiration traits as defined in the main text, where ETS (circles and solid lines) represents the maximal FCCP-stimulated respiration, ROUTINE (squares and dotted-dashed lines) is the unmanipulated respiration on endogenous substrates, OXPHOS (triangles and long-dashed lines) is the respiration directed towards oxidative phosphorylation whereby ATP is produced, and LEAK (rhombuses and short-dashed lines) is the respiration used to offset proton slippage through the inner mitochondrial membrane. Panel B) show flux control ratios (FCR), which are derived as a quotient between two respiration states, specifically: E-R control efficiency (1 - ROUTINE / ETS; circles and solid lines), which is a measure of the proportion of maximum working capacity remaining during endogenous respiration; R-L control efficiency ((ROUTINE - LEAK) / ROUTINE), which is the proportion of endogenous respiration channelled towards ATP production via oxidative phosphorylation (squares and long-dashed lines); E-L coupling efficiency (1 - LEAK / ETS), which is indicative of how 'tightly' electron transport is coupled to ATP production under a stimulated cellular state (triangles and dotted-dashed lines). Respiration traits and FCRs are distinguished by line types and plotting symbols
